# Engineering Oncolytic Measles Virus with MG53 Couples Pyroptotic Tumor Killing with Immune Microenvironment Remodeling to Enhance Checkpoint Immunotherapy in Non-small Cell Lung Cancer

**DOI:** 10.64898/2026.07.15.738733

**Authors:** Zhongguang Li, Cheng Chih Hsu, Fei Jiang, Matthew Bu, Umme Lubaba, Sally L. Li, Serena li Zhao, Pei-Hui Lin, Hunter J. Piegols, Xuefeng Liu, Jianrong Li, Haichang Li

## Abstract

Lung cancer is the leading cause of cancer-related mortality worldwide, with only approximately 30% of patients benefiting from current immune checkpoint immunotherapy. Oncolytic virotherapy offers a promising strategy to overcome this resistance; however, achieving both potent tumor cytotoxicity and robust antitumor immune activation remains a major challenge. Here, we engineered oncolytic measles virus expressing the tumor suppressor MG53 (TRIM72), designated rMeV-MG53, and evaluated its therapeutic potential against non-small cell lung cancer (NSCLC). rMeV-MG53 retained replication kinetics comparable to parental MeV while driving significantly greater cytotoxicity than an unarmed control in NSCLC cells. Mechanistically, MG53 arming amplified caspase-3–dependent apoptosis and potentiated Gasdermin E (GSDME) cleavage, engaging GSDME-mediated pyroptosis as a key tumor-killing mechanism; pharmacological inhibition confirmed caspase-dependent apoptosis as the initiating event leading to pyroptosis, while excluding necroptosis and ferroptosis as significant contributors to rMeV-MG53-induced cytotoxicity. MG53 overexpression induced an intrinsic pro-inflammatory transcriptional signature enriched for TNF, NF-κB, and IL-17 signaling, and rMeV-MG53 elicited significantly stronger type I interferon and pro-inflammatory cytokine induction than the unarmed control. In an immunocompetent, MeV-permissive syngeneic NSCLC model, intratumoral rMeV-MG53 achieved superior tumor growth inhibition, amplified caspase-3/GSDME pyroptosis, selectively upregulated *Cxcl10*, *Ifng*, and *Il1b*, and drove robust CD8⁺ T-cell and Granzyme B⁺ effector infiltration. This immune remodeling was accompanied by compensatory PD-L1 upregulation, and combining rMeV-MG53 with anti-PD-L1 blockade achieved the strongest tumor suppression of all treatment groups. These findings establish MG53-armed oncolytic MeV as a strategy coupling GSDME-mediated pyroptosis with antitumor immune remodeling to sensitize immune-resistant NSCLC to checkpoint immunotherapy.

## Introduction

Lung cancer remains the leading cause of cancer-related mortality worldwide, responsible for approximately 2.5 million new diagnoses and 1.8 million deaths annually as of 2022(1). Non-small cell lung cancer (NSCLC) accounts for approximately 85% of all lung cancer cases, encompassing adenocarcinoma, squamous cell carcinoma, and large cell carcinoma as major histological subtypes(2). Despite considerable advances in targeted therapy and immunotherapy, the five-year overall survival rate for lung cancer remains approximately 25%, underscoring the persistent need for novel therapeutic strategies(3). The advent of immunotherapy such as immune checkpoint inhibitors (ICIs) has revolutionized the treatment landscape for advanced NSCLC, with response rates of 10% -25%(4, 5). Patients responsive to ICI commonly harbor an immune-infiltrated (hot) tumor microenvironment (TME), characterized by pre-existing anti-tumor immunity and abundant immune cell infiltration. However, the remaining 75% - 90% of patients do not respond owing to the immunosuppressive properties (cold) of TME characterized by limited immune cell infiltration into the TME(6–8). Consequently, developing safe and efficacious strategies to reprogram the immunosuppressive TME and enhance anti-tumor immune responses remains a critical unmet medical need.

MG53 (also known as TRIM72) is a member of the tripartite motif (TRIM) protein family that was originally characterized as a key molecule in cell membrane repair(9). Our group and others have extensively demonstrated that recombinant human MG53 (rhMG53) protein exerts broad therapeutic activity across multiple organ systems, including muscular dystrophy(9, 10), lung injury(11, 12), myocardial infarction(13), kidney injury(14, 15), and wound healing(16, 17). Beyond its membrane repair function, accumulating evidence has established MG53 as a critical regulator of cellular homeostasis by modulating oxidative stress, autophagy, and inflammation(18, 19). Importantly, we recently uncovered an antitumor function of MG53 in lung cancer(20). We also demonstrated that rhMG53 acts synergistically with chemotherapeutic agents to promote death of lung cancer cells and suppress tumor growth of drug-resistant cancers in colorectal carcinoma(21). Several subsequent studies further support the function for MG53 as an antitumor therapeutic agent(22–24). The dual properties—tumor suppression and immune modulation—make MG53 an attractive candidate payload for arming an oncolytic virus, potentially coupling enhanced cytotoxic with a more immunogenic mode of tumor cell death.

Oncolytic virotherapy (OV) has emerged as a safe and promising platform in cancer immunotherapy, distinguished by its capacity to selectively lyse tumor cells while sparing normal cells, and to simultaneously reprogram the antitumor immune response state(25–27). The T-Vec (Talimogene laherparepvec) is an attenuated herpes simplex virus type 1 (HSV-1) engineered to encode granulocyte-macrophage colony-stimulating factor (GM-CSF). It received FDA approval in the USA for intratumoral treatment of advanced melanoma and demonstrated synergistic antitumor efficacy when combined with immune checkpoint inhibitors(28). Measles virus (MeV) is a paramyxovirus and another attractive oncolytic virus. Using reverse genetics system, MeV can be engineered to express therapeutic transgenes insertion(27). MeV exhibits inherent tumor tropism mediated by the overexpression of CD46, a complement regulatory protein uniformly upregulated on human tumor cells relative to normal tissues(29, 30). Attenuated vaccine strain of MeV is an attractive oncolytic backbone due to its established clinical safety record, natural tropism for tumor cells overexpressing the viral entry receptor CD46, and capacity to induce immunogenic cell death. Critically, engineering MeV to express therapeutic transgenes— a strategy already validated for cytokines, suicide genes, and immune-stimulatory payloads—offers a modular approach to enhance oncolytic potency and reshape the tumor immune microenvironment. However, the optimal transgene cargo for maximizing both direct cytotoxicity and immunogenicity in NSCLC remains an active area of investigation.

A mechanistically critical modality by which oncolytic viruses potentiate antitumor immunity is through the induction of Gasdermin-mediated pyroptosis(31). Unlike classical apoptosis, which is generally considered immunologically silent, pyroptosis is a lytic, inflammatory form of cell death characterized by membrane pore formation, release of damage-associated molecular patterns (DAMPs), and robust recruitment of immune effector cells(32, 33). GSDME, a key member of the Gasdermin protein family, is specifically cleaved and activated by caspase-3, thereby converting conventional apoptotic signaling into immunogenic pyroptotic cell death(32). GSDME-mediated pyroptosis has been demonstrated to profoundly reshape the TME, converting immunologically "cold" tumors into "hot," immune-infiltrated states and potentiating the efficacy of immune checkpoint blockade across multiple cancer models(33). The caspase-3/GSDME pyroptosis axis thus represents an attractive target for enhancing antitumor immunotherapeutic efficacy.

In this study, we engineered oncolytic MeV expressing antitumor MG53 (rMeV-MG53) and evaluated its therapeutic potential against NSCLC. We hypothesized that MG53 arming would not only preserve the oncolytic activity of MeV but also potentiate caspase-3/GSDME-dependent pyroptosis, thereby converting tumor cell death into a more immunogenic event capable of remodeling the TME. Using a panel of human NSCLC cell lines and a novel immunocompetent, MeV-permissive syngeneic mouse model (LL/2-hCD46), we demonstrate that rMeV-MG53 enhances tumor cell killing through coordinated activation of apoptosis and GSDME-mediated pyroptosis, amplifies antiviral and pro-inflammatory cytokine responses, and drives strong CD8⁺ T-cell and Granzyme B⁺ infiltration into tumors *in vivo*. We further show that this immune remodeling is accompanied by compensatory PD-L1 upregulation, and that combining rMeV-MG53 with anti-PD-L1 blockade achieves superior tumor control compared to either monotherapy. Together, these findings establish MG53 as a rationally designed oncolytic payload that couples enhanced direct cytotoxicity with immune sensitization, supporting a combinatorial strategy to overcome resistance to checkpoint immunotherapy in NSCLC.

## Materials and Methods

### Chemicals, Cells, and Cell Culture

All chemicals were obtained from Sigma-Aldrich unless otherwise indicated. Human cell line A549, H460, H1299, H1975, HEK293T, HEp-2, African green monkey Vero CCL-81, and Murine Lewis lung carcinoma LL/2 cells were purchased from ATCC and maintained at 37°C in a humidified 5% CO₂ incubator. All cells were cultured in RPMI-1640 or DMEM supplemented with 10% fetal bovine serum (FBS), 100 U/ml penicillin and 100 μg/ml streptomycin.

### Engineering and Recovery of Recombinant Oncolytic MeV

All measles viruses used in this study are derived from the attenuated oncolytic MeV Edmonston vaccine strain(34). Plasmids encoding the full-length antigenomic cDNA of MeV with insertion of MG53 or mCherry at the P/M junction was constructed using a yeast-based recombination system described previously(34–38). The *MG53* gene was synthesized by GenScript (Piscataway, NJ). In the yeast-derived plasmids, the insertion regions were amplified by PCR. Positive plasmids were subsequently transformed into DH10B *E. coli* for further amplification. Plasmid DNA was initially screened by restriction enzyme (HindIII) digestion and confirmed by full-length plasmid DNA sequencing to ensure that no additional mutations were introduced during the process. Recovery of recombinant MeV (rMeV) from the infectious cDNA clones was described previously(35, 36). Briefly, HEp-2 cells were infected with recombinant modified vaccinia Ankara virus (MVA-T7) expressing T7 RNA polymerase at 37°C for 1 hour. The plasmid encoding the full-length antigenomic cDNA of MeV with insertion of the MG53 or mCherry was co-transfected with plasmids expressing MeV nucleocapsid complex (pN, pL, and pP) using Lipofectamine 3000 (Thermo Fisher Scientific, Cat. No. L3000015). Four days after transfection, HEp-2 cells were scraped from the plates with a cell scraper and transferred to Vero CCL-81 cells at 90% confluence and co-cultured for 4-7 days to amplify the recombinant virus. Syncytia formation-the characteristic cytopathic effects (CPE) of MeV infection, was observed following co-culture, confirming successful viral recovery. A plaque assay was subsequently performed, and individual plaques were picked and seeded onto fresh Vero CCL-81 cells to eliminate the helper virus (MVA-T7). Purified viruses were titrated by plaque assay on Vero CCL-81 cells and stored at −80°C until use. The resulting recombinant viruses were designated rMeV-MG53 and rMeV-mCherry.

### Preparation of Concentrated rMeV Stocks

Eight T175 flasks of Vero CCL-81 cells were grown to approximately 95% confluency and then infected with indicated recombinant MeV at a multiplicity of infection (MOI) of 0.1. When substantial CPE was observed on days 2 or 3 post-infection, viral supernatant and cell lysates were collected using a cell scraper and centrifuged at 300 × *g* for 5 min at 4°C. The supernatant was collected and kept on ice, while cell pellets were subjected to three freeze-thaw cycles in fresh DMEM containing 10% trehalose, followed by centrifugation at 300 × *g* for 5 min at 4°C. The two supernatant fractions were then combined. A 20% sucrose cushion was prepared in 1× NTE buffer. For ultracentrifugation, 20–25 mL of virus-containing supernatant was carefully loaded into ultracentrifuge tubes, and 5 mL of 20% sucrose solution was gently layered beneath the sample using a syringe fitted with a long needle. The virus was pelleted through the sucrose cushion by ultracentrifugation at 25,000 rpm for 2 hours at 4°C. Following centrifugation, the supernatant was discarded, and the viral pellet was resuspended in 200 μL of 10% trehalose by gentle pipetting. The concentrated virus stocks were aliquoted and stored at −80°C until titration.

### MeV Plaque Assay

Vero CCL-81 cells in 12-well plates were used for infection with recombinant MeV. The virus was serially diluted 10-fold and adsorbed onto the cells for 1 hour at 37°C. Cells were then overlaid with 1 mL of DMEM containing 0.25% (w/v) low-melting agarose, 0.12% (w/v) NaHCO_3_, 2% (w/v) FBS, 25 mM HEPES, 2 mM L-Glutamine, 100 U/ml penicillin, and 100 µg/ml of streptomycin, and incubated at 37°C for 4 days. Cells were then fixed with 4% paraformaldehyde for 1 hour, then agarose overlay was removed, and viral plaques were visualized by staining with 0.05% (w/v) crystal violet in 10% ethanol.

### MeV Growth Curve Assay

Confluent Vero CCL-81 cells in 12-well plates were inoculated with each recombinant virus at an MOI of 0.1. After 1 hour of infection, the inoculum was removed, the cells were washed twice with DMEM, and 1 mL of fresh DMEM with 2% FBS was added. Infected cells were grown at 32℃ incubators with 5% CO_2_. At the indicated time points, cell culture supernatant was collected, and the cell lysates was subjected to three freeze-thaw cycles. The supernatant and cell lysates were then combined, clarified by centrifugation at 300 × *g* for 2 minutes, and the resulting supernatant was used for virus titration by plaque assay on Vero CCL-81 cells.

### Generation of Stable LL/2-hCD46 Cells

To establish the stable LL/2-hCD46 cell line, the hCD46-T2A-GFP fragment (Addgene #205457) was directionally assembled into the PCR-linearized, GFP-excised pLenti-CMV-Puro backbone (Addgene #17448) using the In-Fusion Cloning Kit (Takara Bio, Cat. No. 638951). Following sequence verification, the recombinant plasmid was co-transfected with the lentiviral packaging plasmid psPAX2 and envelope plasmid pMD2.G into HEK293T cells using Lipofectamine 3000 to produce lentiviral particles. Viral supernatants were harvested at 48 and 72 hours post-transfection, pooled, and clarified by centrifugation at 300 × *g* for 10 minutes to remove cellular debris. LL/2 cells were then transduced with the clarified viral supernatant in the presence of 8 μg/mL polybrene (Sigma-Aldrich, Cat. No. TR-1003-50UL). At 48 hours post-infection (48 hpi), stably transduced cells were selected using 2 μg/mL puromycin (Thermo Fisher Scientific, Cat. No. A1113803) until all non-transduced cells had been eliminated. Stable integration and expression of hCD46 and GFP were subsequently confirmed by fluorescence microscopy and Western blot analysis.

### Cell Viability Assay

Cell viability was assessed using the Cell Counting Kit-8 (CCK-8; Dojindo Molecular Technologies, Cat. No. CK04) according to the manufacturer’s instructions. Briefly, cells were seeded into 96-well plates at a density of 2×10^4^ cells per well. After 24 hours of incubation, cells were infected with the rMeV-mCherry or rMeV-MG53 at an MOI of 1.0 for 24h, 48h and 72h. For pathway-dependency experiments, cells were infected at an MOI of 1.0 for 24 h post-infection, and subsequently treated with DMSO (vehicle control), the pan-caspase inhibitor Z-VAD-FMK (Z-VAD, 20 µM), the necroptosis inhibitor necrostatin-1 (Nec-1, 20 µM), or the ferroptosis inhibitor ferrostatin-1 (Fer-1, 10 µM), all from MedChemExpress; and cell viability was then assessed at 48 hpi. Absorbance was measured at 450 nm using a microplate reader, and viability was normalized to mock-treated controls.

### Western Blot Analysis

Cells and tumor tissues were lysed on ice for 20 minutes in RIPA buffer supplemented with protease and phosphatase inhibitors. Following centrifugation (15,000 × *g*, 10 min, 4°C), protein concentrations were determined by bicinchoninic acid assay (Thermo Fisher Scientific, Cat. No. 23225). Equal amounts of protein were denatured at 100°C for 5 minutes in standard SDS sample loading buffer, resolved by SDS-PAGE, and transferred to PVDF membranes. Membranes were blocked with 5% non-fat dry milk in Tris-buffered saline containing 0.1% Tween-20 (TBST) for 1 hour at room temperature then incubated overnight at 4°C with the following primary antibodies: anti-cleaved caspase-3 (Cell Signaling Technology, Cat. No. 9661S), anti-GSDME (Abcam, Cat. No. ab215191), anti-GAPDH (Proteintech, Cat. No. 60004-1-Ig), anti-CD46 (Proteintech, Cat No. 12494-1-AP), and a homemade anti-MG53 antibody(9). After three washes with TBST, membranes were incubated with HRP-conjugated anti-rabbit IgG secondary antibody (Cell Signaling Technology, Cat. No. 7074) for 1 hour at room temperature. Protein bands were visualized using enhanced chemiluminescence (ECL) substrate (Thermo Fisher Scientific, Cat. No. 34095) and quantified using ImageJ software (NIH, Bethesda, MD, USA; version 1.54p).

### Immunofluorescence Staining (IF)

Cells were washed with PBS, fixed with 4% paraformaldehyde for 10 min, and permeabilized with 0.5% Triton X-100 for 5 min. Following blocking in 3% bovine serum albumin (BSA) for 1 hour at room temperature, cells were incubated with anti-MG53 antibody (1:400) overnight at 4°C. Cells were then washed three times with PBS and incubated with an Alexa Fluor 488-conjugated goat anti-rabbit secondary antibody (1:500; Thermo Fisher Scientific, Cat. No. A-11008) for 1 hour at room temperature. Nuclei were counterstained with DAPI, and coverslips were mounted using an antifade mounting medium. Images were captured and analyzed using a Nikon ECLIPSE Ti2 confocal microscope.

### RNA Sequencing and Transcriptomic Analysis

Total RNA was extracted from A549 and H460 cells stably overexpressing MG53 and their respective empty vector controls using TRIzol reagent (Invitrogen, Cat. No. 15596026) according to the manufacturer’s instructions. RNA quantity and integrity were assessed, and the qualified samples were submitted to Innomics Inc. for library preparation, high-throughput sequencing, and bioinformatic analysis. Briefly, mRNA was enriched from total RNA using oligo(dT) magnetic beads, fragmented, and reverse-transcribed into cDNA to construct sequencing libraries. Libraries were then sequenced on an Illumina NovaSeq platform to generate paired-end reads.

For bioinformatic analysis, raw reads were filtered to remove adapters and low-quality sequences. The resulting clean reads were mapped to the human reference genome using HISAT2 (v2.2.1). Read counts for each gene were quantified, and differential expression analysis between MG53-overexpression and control groups was performed using the DESeq2 (v1.4.5) R package. Genes meeting the criteria of an adjusted *p*-value ≤ 0.05 and absolute log₂ fold change > 1 were defined as significantly differentially expressed genes (DEGs). Functional enrichment analysis of DEGs, including Gene Ontology (GO) and Kyoto Encyclopedia of Genes and Genomes (KEGG) pathways analysis, was subsequently performed to determine the primary biological functions and pathways regulated by MG53 overexpression. Results were visualized as bubble plots, in which bubble size represents the number of enriched genes within each pathway and color intensity reflects the adjusted *p*-value (Q-value).

### Reverse Transcription Quantitative Real-Time PCR (RT-qPCR)

H460 cells were infected with rMeV-mCherry or rMeV-MG53 at MOIs of 1.0 and 2.0 for 48 hours. Total RNA from cells and mouse tumor tissues was isolated using TRIzol reagent (Invitrogen, Cat. No. 15596026) according to the manufacturer’s instructions. 1µg of total RNA was reverse-transcribed into cDNA using the Radiant™ cDNA Synthesis Kit, 1-Step (Alkali Scientific, Cat. No. QC1155). RT-qPCR was performed using the Radiant™ Green qPCR Master Mix (Alkali Scientific, Cat No. QS1005) on a QuantStudio 3 Real-Time PCR System (Applied Biosystems, Thermo Fisher Scientific). Relative mRNA expression levels of *IFNB1*, *IL6,* and *IL1B* (in H460 cells) and *Cxcl10*, *Ifng*, *Tnf*, and *Il1b* (in mouse tumors) were quantified using the 2^−ΔΔCt^ method and normalized to *GAPDH* (human) or *Actb* (mouse) as reference genes. Primer sequences are listed in **Table 1**.

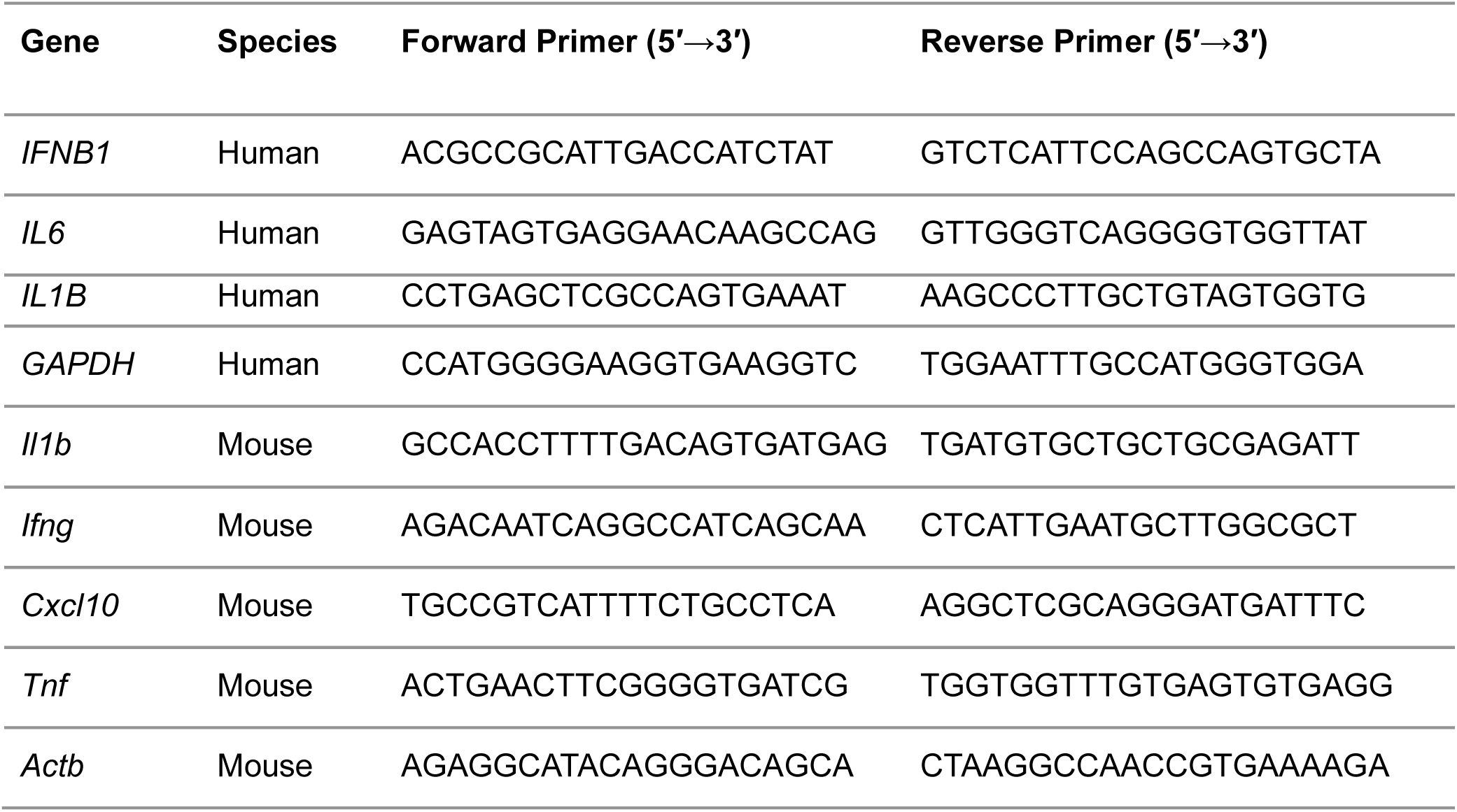

### Syngeneic Tumor Models and *In Vivo* Treatment

All animal care and use procedures were performed in accordance with NIH guidelines and approval by the Institutional Animal Care and Use Committee (IACUC) of The Ohio State University. *MG53* knockout (KO, *MG53^-/-^*) mice and their wild type littermates were bred and generated as previously described(9).

To evaluate the tumor-suppressive role of endogenous MG53, C57BL/6 wild-type (WT) and *MG53^-/-^* mice (8-10 weeks old) were subcutaneously inoculated with 5×10^5^ LL/2 cells into the right flank. Tumor volumes were measured every other day using digital calipers and calculated using the standard formula: Tumor size = (length × width²) / 2. On day 16 post-implantation, mice were humanely euthanized, and tumors were excised and weighed.

For the monotherapy efficacy cohort (**Figure 4F–I**), LL/2-hCD46 cells (5 × 10⁵ cells) were subcutaneously implanted into the right flank of C57BL/6 WT mice. When tumors reached a palpable size on day 7 post-implantation, mice were randomly divided into three groups (mock, rMeV-mCherry, and rMeV-MG53) and received intratumoral injections of recombinant virus (1 × 10⁶ plaque-forming units [PFU] in 100 µL per dose) once daily on days 7–14 post-implantation; tumor growth was monitored to day 15. For the combination cohort (**Figure 6**), performed independently, established LL/2-hCD46 tumors were randomly assigned to five groups—mock, anti-PD-L1, rMeV-mCherry, rMeV-MG53, and rMeV-MG53 plus anti-PD-L1 (*n* =6-10 per group). Recombinant virus (1 × 10⁶ PFU in 100 µL per dose) and anti-PD-L1 antibody (clone 10F.9G2, Bio X Cell, Lebanon, NH, USA; 5 mg/kg) were co-administered intratumorally once daily on days 7–14 post-implantation. In both cohorts, tumor volume was measured every other day with digital calipers and calculated as (length × width²)/2.

At the end of the experiment, all mice were euthanized. Tumors were excised and divided into two portions: one was snap-frozen at −80°C for protein and RNA extraction, and the other was fixed in 4% paraformaldehyde for 24 hours and processed for paraffin embedding and sectioning. Major organs including heart, liver, spleen, lungs, kidneys, and brain, were also collected, fixed in 4% paraformaldehyde for 24 hours, and subjected to hematoxylin and eosin (H&E) staining to assess potential systemic toxicity.

### Histological Staining and Immunohistochemistry (IHC)

Histology staining and immunohistochemistry (IHC) were performed as previously described(16). Briefly, tissue samples were fixed in 4% paraformaldehyde for 24 hours, dehydrated, embedded in paraffin, and sectioned at 5 μm thickness. Prior to staining, all sections were deparaffinized in xylene and rehydrated through a graded ethanol series. For histological evaluation, sections were routinely stained with H&E. For IHC staining, rehydrated sections underwent heat-induced antigen retrieval in EDTA buffer (pH 9.0) for 20 minutes. Endogenous peroxidase activity was quenched with 3% H_2_O_2_ for 10 minutes, followed by blocking with 5% BSA in PBS for 1 hour at room temperature. Sections were then incubated overnight at 4°C with the following primary antibodies (All from Cell Signaling Technology, 1:300 dilution): anti-CD8a (Cat. No. 98941S), anti-Ki67 (Cat. No. 12202S), anti-Granzyme B (Cat No. 44153S), and anti-PD-L1 (Cat. No. 80517S). After washing with PBST, slides were incubated with the SignalStain® Boost IHC Detection Reagent (HRP, Rabbit; Cell Signaling Technology, Cat. No. 8114P) for 30 minutes at room temperature. Immunoreactivity was visualized using a DAB substrate kit (Cell Signaling Technology, Cat. No. 8059S), and nuclei were counterstained with hematoxylin. All H&E and IHC slides were dehydrated, cleared in xylene, coverslip-mounted, and imaged using an Olympus BX43 upright microscope. Quantitative analysis of the images was performed using ImageJ software (NIH, Bethesda, MD, USA; version 1.54p).

### Statistical Analysis

All data were analyzed using GraphPad Prism (version 10.0, GraphPad Software, La Jolla, CA, USA) and are presented as mean ± standard deviation (SD). For comparisons between two groups, statistical significance was determined using unpaired two-tailed Student’s *t*-test. For multiple group comparisons, one-way or two-way ANOVA followed by Tukey’s multiple-comparisons post-hoc test or Dunnett’s T3 post hoc test was applied as appropriate. A *p-*value < 0.05 was considered statistically significant. Statistical significance is denoted as \**p* < 0.05; \*\**p* < 0.01; \*\*\**p* < 0.001; and \*\*\*\**p* < 0.0001; ns, not significant.

## Results

### Recovery and characterization of recombinant measles virus expressing full-length MG53 (rMeV-MG53)

MG53 (TRIM72) contains conserved RING, B-box, coiled-coil, and C-terminal PRY-SPRY domains (**Figure 1A**). MeV is a non-segmented negative-sense RNA virus encoding six genes in the order 3′-Leader-N-P-M-F-H-L-Trailer 5′ (**Figure 1B**). First, we engineered MeV to express tumor-suppressive MG53. We chose to insert MG53 gene into the junction between the P and M genes in MeV genome because foreign gene inserted at this position achieved high protein expression without significantly disturbing virus replication(34, 37). Briefly, the full-length human MG53 coding sequence or mCherry gene was amplified by PCR and inserted into the junction between P and M genes in the genome of the attenuated MeV Edmonston vaccine strain using a yeast-based recombination system(38, 39). Using reverse genetic system, recombinant MeV expressing MG53 or mCherry was rescued from infectious cDNA clones, plaque purified, and designated rMeV-MG53 and rMeV-mCherry, respectively. Subsequently, the entire genome of each recombinant virus was sequenced to confirm that it contained the desired gene insertion, and no additional mutations had been introduced. Plaque assay showed that plaque morphology formed by these two recombinant viruses was comparable to the parental wild-type virus (MeV-WT) (**Figure 1C**), suggesting that insertion of MG53 or mCherry does not significantly disturb virus spread.

**Figure 1.**
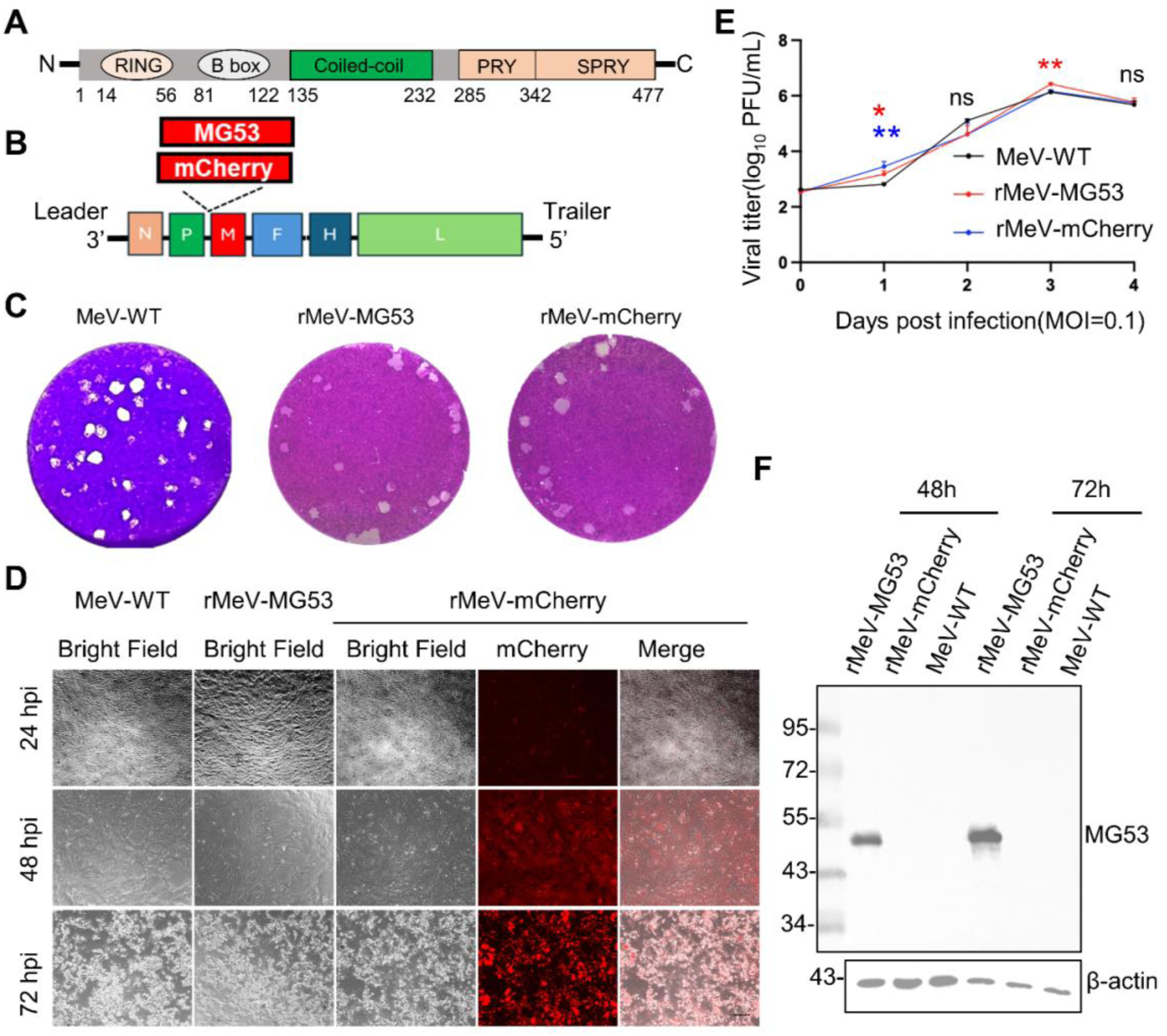
Recovery and characterization of rMeV-MG53, an MG53-armed recombinant oncolytic measles virus. **(A)** Schematic representation of the human MG53 (TRIM72) protein domain structure, showing the RING, B-box, Coiled-Coil, PRY, and SPRY domains. **(B)** Genome organization of MeV Edmonston vaccine strain. The MG53 or mCherry transcription unit was inserted into the junction between the phosphoprotein (P) and matrix (M) genes in MeV genome to generate rMeV-MG53 and rMeV-mCherry, respectively. **(C)** Representative plaque morphology of MeV-WT, rMeV-MG53, and rMeV-mCherry in Vero CCL-81 cells. Plaques were developed at day 4 after virus infection and visualized by crystal violet staining. **(D)** Representative bright-field and fluorescence images showing CPE induced by MeV-WT, rMeV-MG53, and rMeV-mCherry in Vero CCL-81 cells at 24, 48, and 72 h post-infection (hpi). mCherry fluorescence confirms transgene expression from the recombinant virus. Scale bar, 100 μm. **(E)** Multi-step growth kinetics of MeV-WT, rMeV-MG53, and rMeV-mCherry in Vero CCL-81 cells. Cells were infected at an MOI of 0.1, and viral titers (log₁₀ PFU/mL) were determined by plaque assay. Data are presented as mean ± SD from three independent experiments. Statistical significance was determined by two-way ANOVA with Geisser–Greenhouse correction followed by Tukey’s multiple-comparisons test. Red asterisks indicate comparisons between rMeV-MG53 and MeV-WT, blue asterisks indicate comparisons between rMeV-mCherry and MeV-WT. \**p* < 0.05, \*\**p* < 0.01; ns, not significant. **(F)** Immunoblot analysis confirming MG53 expression. Vero CCL-81 cells were infected with rMeV-MG53, rMeV-mCherry, or MeV-WT at an MOI of 1.0, and cell lysates were collected at 48 and 72 hpi for Western blot. β-actin served as a loading control

To evaluate replication kinetics, Vero CCL-81 cells were infected with MeV-WT, rMeV-MG53, or rMeV-mCherry, cytopathic effects (CPE) were monitored, and cell culture supernatant and lysate were collected for virus titration. Similar to MeV-WT and rMeV-mCherry, rMeV-MG53 productively infected Vero CCL-81 cells, inducing characteristic CPE including syncytia formation that expanded progressively at 24, 48, and 72 h post-infection (**Figure 1D**). Multi-step growth curves showed that all three viruses replicated to comparably high titers and reached peak titers at 72 h post-infection (**Figure 1E**). Although rMeV-MG53 and rMeV-mCherry showed modestly lower titers than MeV-WT at early time points (day 1, *p* < 0.05 and *p* < 0.01, respectively) and rMeV-MG53 a transient difference at day 3 (*p* < 0.01), the three viruses converged to equivalent peak titers by 72 h, with no significant differences at later time points (days 2 and 4; ns). These results indicate that insertion of MG53 or mCherry causes at most a minor, transient kinetic lag without compromising overall replication competence or peak viral yield.

To confirm MG53 protein expression, Vero CCL-81 cells were infected with MeV-WT, rMeV-MG53, or rMeV-mCherry, and MG53 protein was assessed by Western blot. Robust MG53 protein was detected in rMeV-MG53–infected cells but not in the cells infected with rMeV-mCherry or MeV-WT (**Figure 1F**). These data demonstrate that MG53 is highly expressed by MeV vector.

### rMeV-MG53 induces significant cell death in NSCLC cells through activation of apoptosis and GSDME-mediated pyroptosis

We next evaluated the oncolytic activity of rMeV-MG53 against human lung cancer cells. A panel of NSCLC cell lines (A549, H460, H1299, and H1975) were infected with rMeV-mCherry or rMeV-MG53. Infection in the rMeV-mCherry group was tracked by mCherry fluorescence, whereas productive rMeV-MG53 infection was confirmed by MG53 immunofluorescence together with the progression of CPE (syncytia formation) of MeV replication. Productive viral infection and replication were confirmed across all tested NSCLC cell lines, as evidenced by widespread mCherry fluorescence signals detected as early as 24 hpi. Over the course of infection, mCherry fluorescence intensity increased dramatically, culminating in extensive cell-cell fusion and syncytia formation followed by massive cell death at 72 hpi — hallmarks of productive oncolytic MeV replication (**Figure 2A**). Successful MG53 delivery was verified by immunofluorescence, which showed strong MG53 signals in rMeV-MG53–infected cells (**Figure 2B**), and by Western blotting confirming MG53 expression across all four lines (**Figure 2C**). Importantly, rMeV-MG53 reduced cell viability more effectively than the vector control (rMeV-mCherry). While cell viability remained comparable at 24 hpi, the rMeV-MG53 virus showed significantly lower viability than rMeV-mCherry in all four lines by 48 hpi (A549, *p* < 0.001; H460, *p* < 0.001; H1299, *p* < 0.01; H1975, *p* < 0.001). This enhanced oncolytic activity persisted at 72 hpi in A549 (*p* < 0.05) and H1975 (*p* < 0.001), whereas H460 and H1299 cells reached similarly low viability in both virus groups (**Figure 2D**). In summary, MG53 arming preserves MeV infectivity and enhances its oncolytic activity across a diverse panel of NSCLC cell lines.

**Figure 2.**
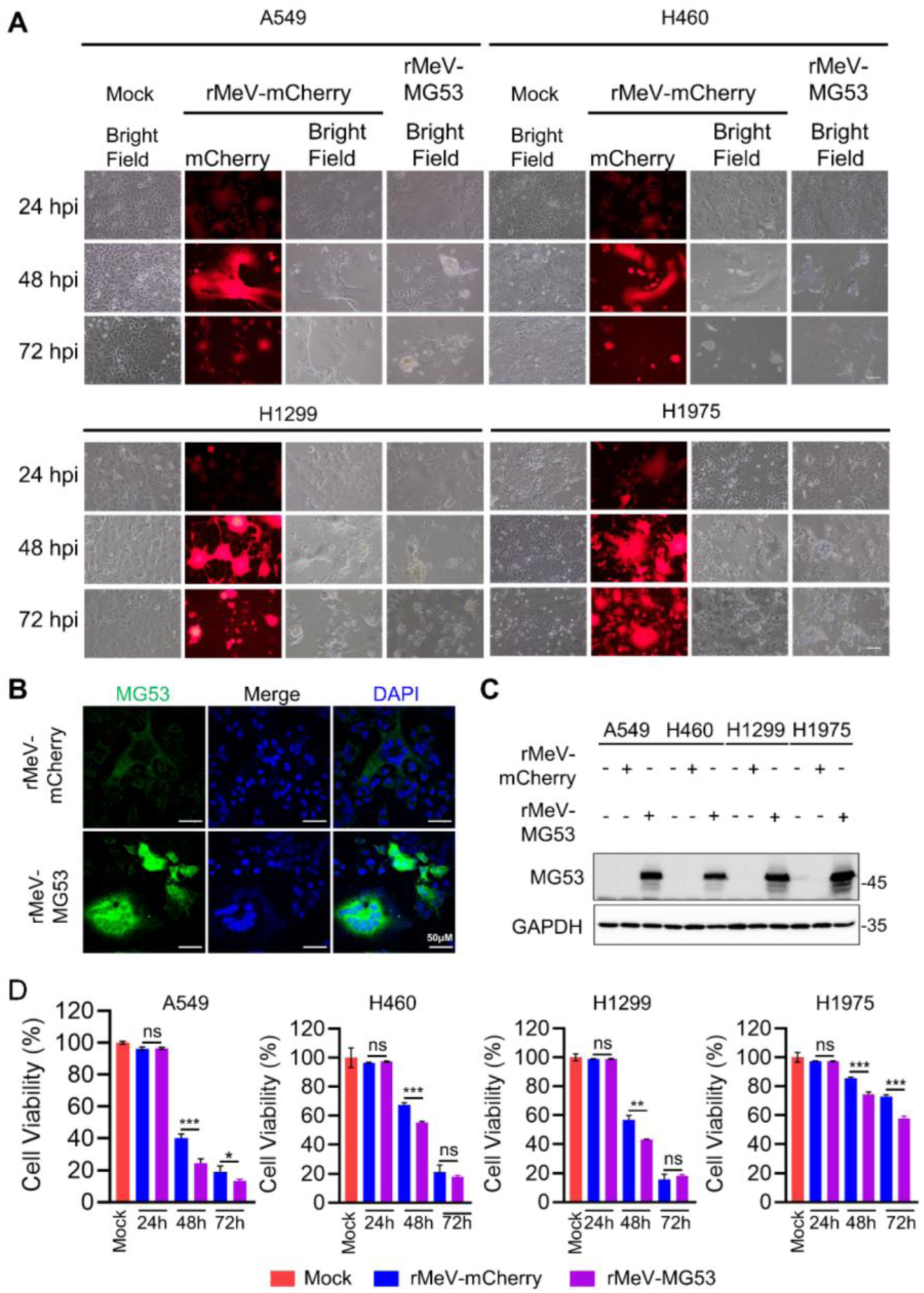
rMeV-MG53 productively infects NSCLC cells and delivers functional MG53 to enhance oncolytic cytotoxicity. **(A)** Representative bright-field and fluorescence images of A549, H460, H1299, and H1975 cells mock-treated or infected with rMeV-mCherry or rMeV-MG53 at an MOI of 1.0, acquired at 24, 48, and 72 h post-infection (hpi). mCherry fluorescence was used to monitor viral infection and replication. Scale bar, 50 μm. **(B)** Immunofluorescence staining of MG53 (green) in A549 cells infected with rMeV-mCherry or rMeV-MG53 at 24 hpi. Nuclei were counterstained with DAPI (blue). Scale bar, 50 μm. **(C)** Immunoblot analysis of MG53 protein expression in A549, H460, H1299, and H1975 cells infected with rMeV-mCherry or rMeV-MG53 at 24 hpi. GAPDH served as a loading control. **(D)** Cell viability of A549, H460, H1299, and H1975 cells following infection with rMeV-mCherry or rMeV-MG53 at the indicated time points. Cell viability was measured using the CCK-8 assay and normalized to mock-infected controls. Data are presented as mean ± SD (*n*=4 per group). Statistical significance was determined by two-tailed unpaired Student’s *t*-test comparing the rMeV-MG53 and rMeV-mCherry groups at each time point. \**p* < 0.05; \*\**p* < 0.01; \*\*\**p* < 0.001; ns, not significant.

To delineate the mechanistic basis of this enhanced cytotoxicity, we examined the cell death modalities engaged following infection. Phase-contrast microscopy revealed that rMeV-MG53–infected NSCLC cells exhibited prominent cellular swelling and large membrane ballooning at 48 and 72 hpi (**Figure 2A**), the cardinal morphological hallmarks of pyroptosis that are distinct from classical apoptotic morphology(32), suggesting that rMeV-MG53 engages an inflammatory cell death program beyond conventional apoptosis. To characterize the relative contributions of apoptosis and pyroptosis, we assessed key molecular markers of both pathways by Western blot. rMeV-mCherry alone induced apoptosis, as evidenced by detectable cleavage of caspase-3, and rMeV-MG53 infection further increased cleaved caspase-3 levels relative to rMeV-mCherry, indicating that MG53 expression amplifies apoptotic signaling during oncolytic MeV infection (**Figure 3A**). Concurrent assessment of pyroptosis markers showed that both viruses induced cleavage of GSDME, generating the pore-forming N-terminal fragment (GSDME-N). Importantly, GSDME-N was markedly elevated in rMeV-MG53–infected NSCLC cells relative to the rMeV-mCherry–infected controls (**Figure 3A**), indicating that MG53 potentiates the caspase-3/GSDME pyroptotic axis during oncolytic MeV infection. Together, these findings indicate that rMeV-MG53 simultaneously amplifies caspase-3–dependent apoptosis and GSDME-mediated pyroptosis, providing a coherent mechanistic basis for its enhanced cytotoxic activity over the rMeV-mCherry vector control and supporting the caspase-3/GSDME axis as a key driver of rMeV-MG53 antitumor activity in lung cancer.

**Figure 3.**
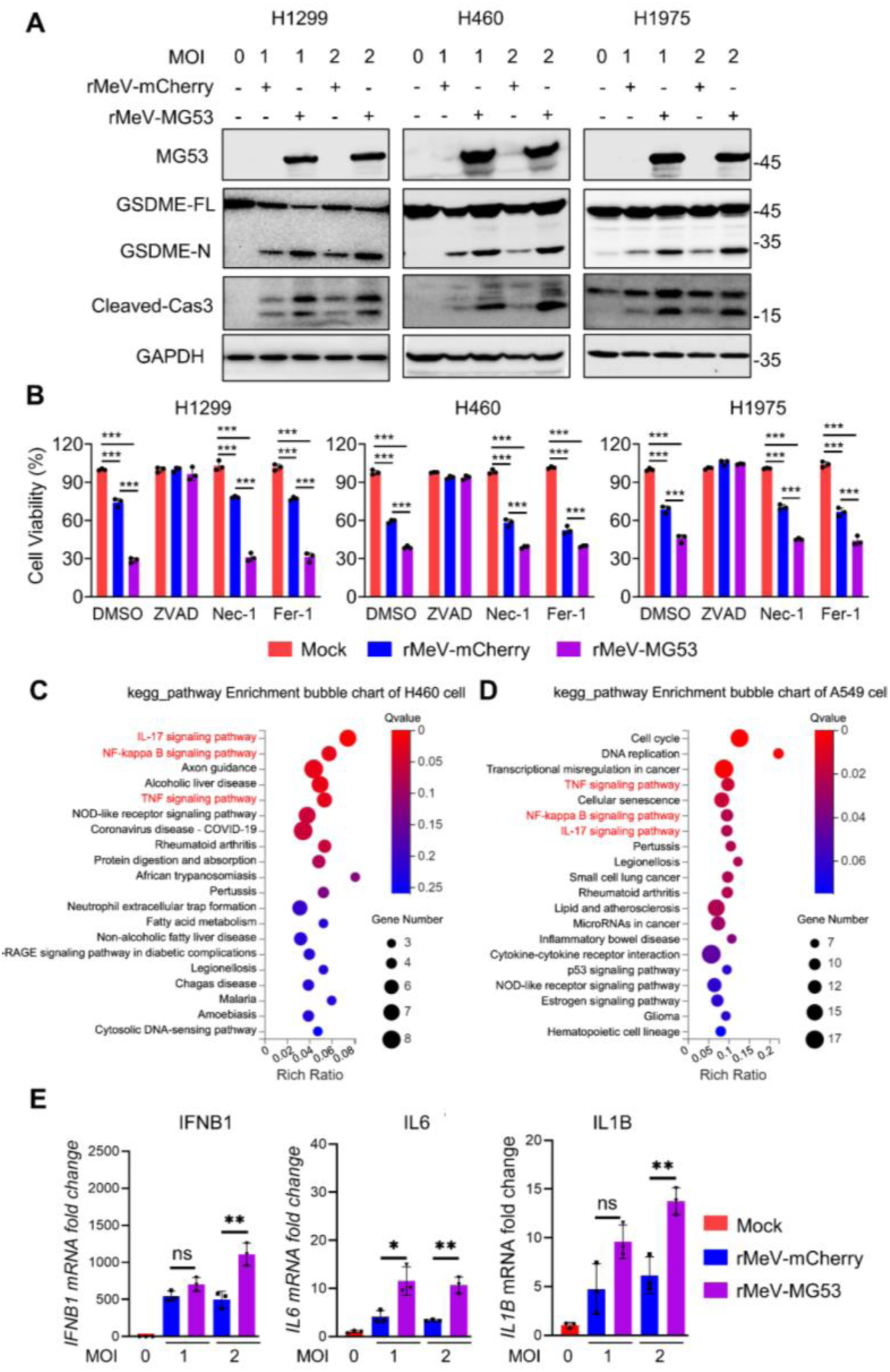
rMeV-MG53 potentiates caspase-3/GSDME-mediated pyroptosis and pro-inflammatory cytokine induction in NSCLC cells. **(A)** Immunoblot analysis of MG53 expression and pyroptosis-related proteins in H1299, H460, and H1975 cells infected with rMeV-mCherry or rMeV-MG53 at the indicated MOI for 48 hpi. Full-length GSDME (GSDME-FL), the cleaved GSDME N-terminal fragment (GSDME-N), and cleaved caspase-3 (Cleaved-Casp3) were examined. GAPDH served as a loading control. (**B)** Cell viability of H1299, H460, and H1975 cells infected with rMeV-mCherry or rMeV-MG53 in the presence of DMSO (vehicle control), Z-VAD-FMK (Z-VAD, pan-caspase inhibitor), necrostatin-1 (Nec-1, necroptosis inhibitor), or ferrostatin-1 (Fer-1, ferroptosis inhibitor) at 48hpi. Data are presented as mean ± SD (*n*=3 per group). \*\*\**p* < 0.001. **(C, D)** KEGG pathway enrichment analysis of differentially expressed genes identified by RNA sequencing in MG53-overexpressing H460 **(C)** and A549 **(D)** cells in the absence of viral infection. Selected immune- and inflammation-related pathways are highlighted in red. Bubble size represents the number of enriched genes; and bubble color indicates adjusted *P* value (Q value). **(E)** RT-qPCR analysis of *IFNB1*, *IL6*, and *IL1B* mRNA expression in H460 cells infected with rMeV-mCherry or rMeV-MG53 at the indicated MOIs for 48 hpi. Gene expression levels were normalized to the internal reference gene (*GAPDH*) and expressed as fold change relative to the mock-infected controls. Data are presented as mean ± SD (*n*=3 per group). Statistical significance was determined by two-tailed unpaired Student’s *t*-test comparing rMeV-MG53 and rMeV-mCherry groups at each MOI. \**p* < 0.05; \*\**p* < 0.01; ns, not significant.

**Figure 4.**
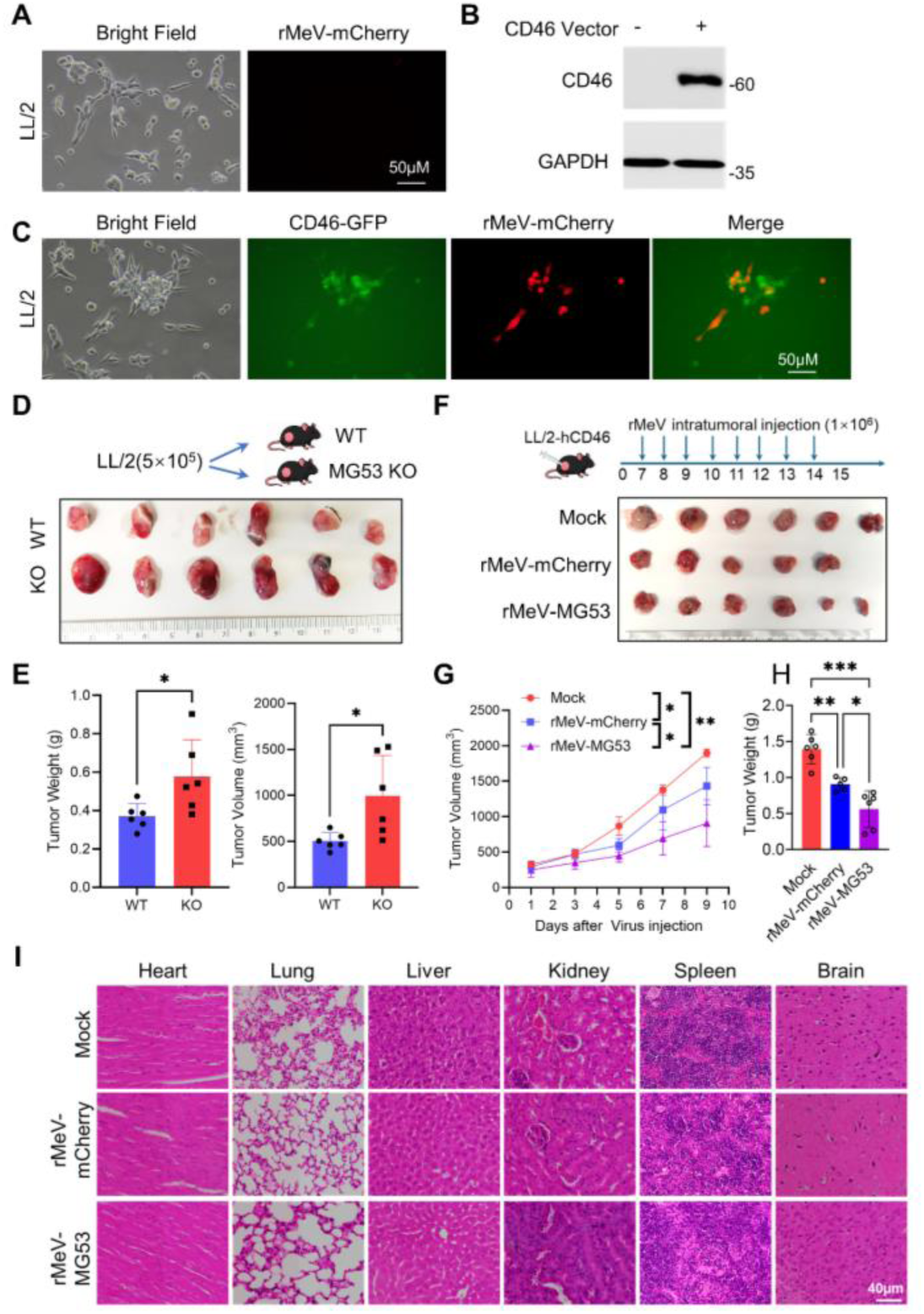
Host MG53 suppresses tumorigenesis and rMeV-MG53 achieves superior tumor control in a MeV-permissive syngeneic NSCLC model. **(A)** Representative bright-field and fluorescence images of parental LL/2 cells following infection with rMeV-mCherry. No mCherry was detected at 72hpi, demonstrating resistance to MeV infection. Scale bar, 50 µm. **(B)** Immunoblot analysis confirming stable expression of hCD46 in LL/2 cells transduced with a hCD46 expressional lentiviral vector. GAPDH served as a loading control. **(C)** Representative bright-field, hCD46-GFP fluorescence, rMeV-mCherry fluorescence, and merged images of LL/2-hCD46 cells following viral infection. Co-localization of mCherry signal with hCD46-GFP-positive cells confirmed efficient MeV entry and replication in hCD46-expressing cells. Scale bar, 50 μm. **(D, E)** LL/2 cells (5 × 10⁵ cells per mouse) were subcutaneously implanted into wild-type (WT) or MG53-knockout (MG53 KO) mice. Representative excised tumors at endpoint **(D)** and quantification of tumor weight and volume **(E)** are shown. Data are presented as mean ± SD (*n* = 6 mice per group). Statistical significance was determined by unpaired two-tailed Student’s *t*-test. \**p* < 0.05. **(F–H)** Therapeutic efficacy of rMeV-MG53 in the LL/2-hCD46 syngeneic tumor model. Mice bearing established LL/2-hCD46 tumors received intratumoral injections of mock, rMeV-mCherry or rMeV-MG53 (1 × 10⁶ PFU per dose, daily from day 7 through day 14 post tumor implantation). Representative excised tumors at endpoint (day 15) **(F)**, tumor growth curves **(G),** and tumor weights **(H)** are shown. **(I)** Representative H&E stainings of heart, lung, liver, kidney, spleen, and brain from virus-treated and mock-treated mice at endpoint, assessing systemic toxicity. Scale bar, 40 µm. Data are presented as mean ± SD (Mock, *n* = 6; rMeV-MG53, *n* = 6; rMeV-mCherry, *n* = 5.). Tumor growth curves were analyzed by two-way repeated-measures ANOVA (treatment × time) with Geisser– Greenhouse correction and Tukey’s multiple-comparisons test. Endpoint tumor weight was analyzed by Brown– Forsythe and Welch ANOVA with Dunnett’s T3 post hoc test. \**p* < 0.05; \*\**p* < 0.01; \*\*\**p* < 0.001.

To pharmacologically define the death pathway engaged, we treated the infected NSCLC cells at 24 hpi with the pan-caspase inhibitor Z-VAD-FMK, the necroptosis inhibitor necrostatin-1 (Nec-1), or the ferroptosis inhibitor ferrostatin-1 (Fer-1), and assessed cell viability at 48 hpi. Only Z-VAD-FMK significantly rescued cell viability in both rMeV-mCherry– and rMeV-MG53–infected cells, while Nec-1 and Fer-1 had no appreciable effects (**Figure 3B**). These results pharmacologically confirm that rMeV-mCherry and rMeV-MG53 induce caspase-dependent cell death as the dominant cytotoxic mechanism and exclude necroptosis and ferroptosis as significant contributors to rMeV-MG53–mediated tumor cell killing.

### MG53 arming engages intrinsic inflammatory programs and amplifies antiviral cytokine output

To determine whether MG53 intrinsically modulates inflammatory signaling independent of viral infection, we profiled the transcriptomes of A549 and H460 cells overexpressing MG53 alone. KEGG pathway enrichment analysis revealed enrichment of immune and inflammatory networks in both cell lines, with TNF α, NF-κB, and IL-17 signaling among the top-ranked pathways (**Figure 3C and D**). Because this transcriptomic signature was generated in the absence of virus infection, it reflects an MG53-intrinsic effect rather than a virus infection-driven response, indicating that MG53 is not transcriptionally inert but instead actively primes cells toward a pro-inflammatory state.

We next investigated whether this intrinsic property translates into amplified cytokine induction during active infection. Briefly, cells were infected by rMeV-MG53 or rMeV-mCherry at MOIs of 1.0 and 2.0, cytokines mRNA expression were quantified by RT-qPCR. We observed that both recombinant viruses induced a high level of expression of type I interferon (*IFNB1*) and pro-inflammatory cytokines (*IL6* and *IL1B*) compared to the mock-infected cells. Notably, rMeV-MG53 elicited a significantly more potent induction than rMeV-mCherry at an MOI of 2.0 across all three cytokines (*IFNB1*, *p* < 0.01; *IL6*, *p* < 0.01; *IL1B*, *p* < 0.01). At an MOI of 1.0, *IL6* induction was also significantly higher in the rMeV-MG53 group (*p* < 0.05), whereas differences for *IFNB1* and *IL1B* were not statistically significant (**Figure 3E**). Collectively, these two independent lines of evidence — an MG53-driven inflammatory transcriptional signature and potentiated induction of antiviral and pro-inflammatory cytokines during infection — demonstrate that MG53 arming reinforces the immunogenic profile of oncolytic MeV.

### Intratumoral rMeV-MG53 enhances oncolytic virotherapy and inhibits tumorigenesis in an immunocompetent lung cancer mouse model

Evaluating oncolytic MeV *in vivo* requires an immunocompetent, virus-permissive model, but murine cells lack human CD46 (hCD46), the entry receptor for MeV. As expected, murine lung cancer cells LL/2 were resistant to rMeV-mCherry infection, showing no fluorescent signal while maintaining normal cell morphology (**Figure 4A**). To establish viral susceptibility, we generated LL/2 cells by expressing hCD46 (LL/2-hCD46) using lentiviral transduction with the hCD46-T2A-GFP plasmid. Ectopic hCD46 expression was confirmed *via* immunoblotting (**Figure 4B**). Crucially, the resulting hCD46-GFP⁺ cells successfully supported viral entry and replication, as evidenced by the robust co-localization of rMeV-mCherry fluorescence within hCD46-GFP-expressing cells (**Figure 4C**). Thus, this engineered LL/2-hCD46 line establishes a critical syngeneic, MeV-permissive platform for evaluating MG53-armed virotherapy within fully immunocompetent hosts.

Before testing therapeutic MG53 delivery, we asked whether endogenous MG53 limits tumor growth in our model. We implanted LL/2 cells (5 × 10⁵ cells) into wild-type (WT) and MG53-knockout (KO) mice. Tumors in MG53-KO mice grew significantly larger and heavier than those in WT mice (tumor weight and volume, both *p* < 0.05) (**Figure 4D and E**). This demonstrates that host MG53 possesses intrinsic tumor-suppressive properties in this NSCLC model, providing a clear biological rationale for using an MG53 armed oncolytic virus to amplify intratumoral MG53. Although this host-derived MG53 effect is mechanistically distinct from the virus-delivered therapy tested below, both support the premise that increasing MG53 in TME suppresses tumorigenesis.

Next, we evaluated therapeutic efficacy in an independent animal cohort. Mice with established LL/2-hCD46 tumors received daily intratumoral injections (1 × 10⁶ PFU in 100 µL per dose, days 7–14) and were monitored until day 15 (**Figure 4F**). Both viruses effectively inhibited tumor growth compared to the mock group, but rMeV-MG53 yielded the most pronounced tumor suppression. Excised tumors were smallest in the rMeV-MG53 group (**Figure 4F**). Tumor-volume curves clearly diverged over time: mock tumors grew the fastest; rMeV-mCherry showed intermediate growth; and rMeV-MG53 grew the slowest (**Figure 4G**). Between-group differences were highly significant (mock vs. rMeV-mCherry, *p* < 0.05; mock vs. rMeV-MG53, *p* < 0.01; rMeV-mCherry vs. rMeV-MG53, *p* < 0.05), indicating that MG53 arming significantly enhances the intrinsic oncolytic capacity of the MeV vector. Endpoint tumor weights mirrored these results, with rMeV-MG53 achieving the lowest tumor burden (mock vs. rMeV-mCherry, *p* < 0.01; mock vs. rMeV-MG53, *p* < 0.001; rMeV-mCherry vs. rMeV-MG53, *p* < 0.05) (**Figure 4H**).

Finally, we assessed systemic tolerability by histologically examining major organs at the endpoint. H&E staining of the heart, lungs, liver, kidneys, spleen, and brain showed no obvious treatment-related pathology in the virus-treated mice relative to the mock controls (**Figure 4I**), suggesting that intratumoral administration restricts virus-mediated effects largely to the tumor site without detectable systemic toxicity, although formal viral biodistribution was not assessed. Overall, these findings indicate that MG53-armed oncolytic MeV achieves greater local tumor control with no apparent systemic toxicity, supporting its use in combination therapies.

### rMeV-MG53 drives *in vivo* pyroptosis and remodels the tumor immune microenvironment

We next investigated whether the pyroptotic and immunogenic mechanisms defined *in vitro* operate within the TME *in vivo*. Immunoblotting of LL/2-hCD46 tumor lysates confirmed that MG53 was exclusively detected in the rMeV-MG53 group. This group also exhibited the most pronounced caspase-3 cleavage and subsequent GSDME-FL to GSDME-N conversion, followed by rMeV-mCherry, with the mock group showing basal levels (**Figure 5A**). Densitometric quantification confirmed that both GSDME-N and cleaved-caspase-3 (normalized to GAPDH) were significantly elevated by either virus compared to mock (*p* < 0.0001), and were further potentiated by MG53 arming relative to the parental rMeV-mCherry (*p* < 0.01) (**Figure 5B**). These data establish that MG53 intrinsically amplifies caspase-3/GSDME-dependent pyroptosis within the solid tumor.

**Figure 5.**
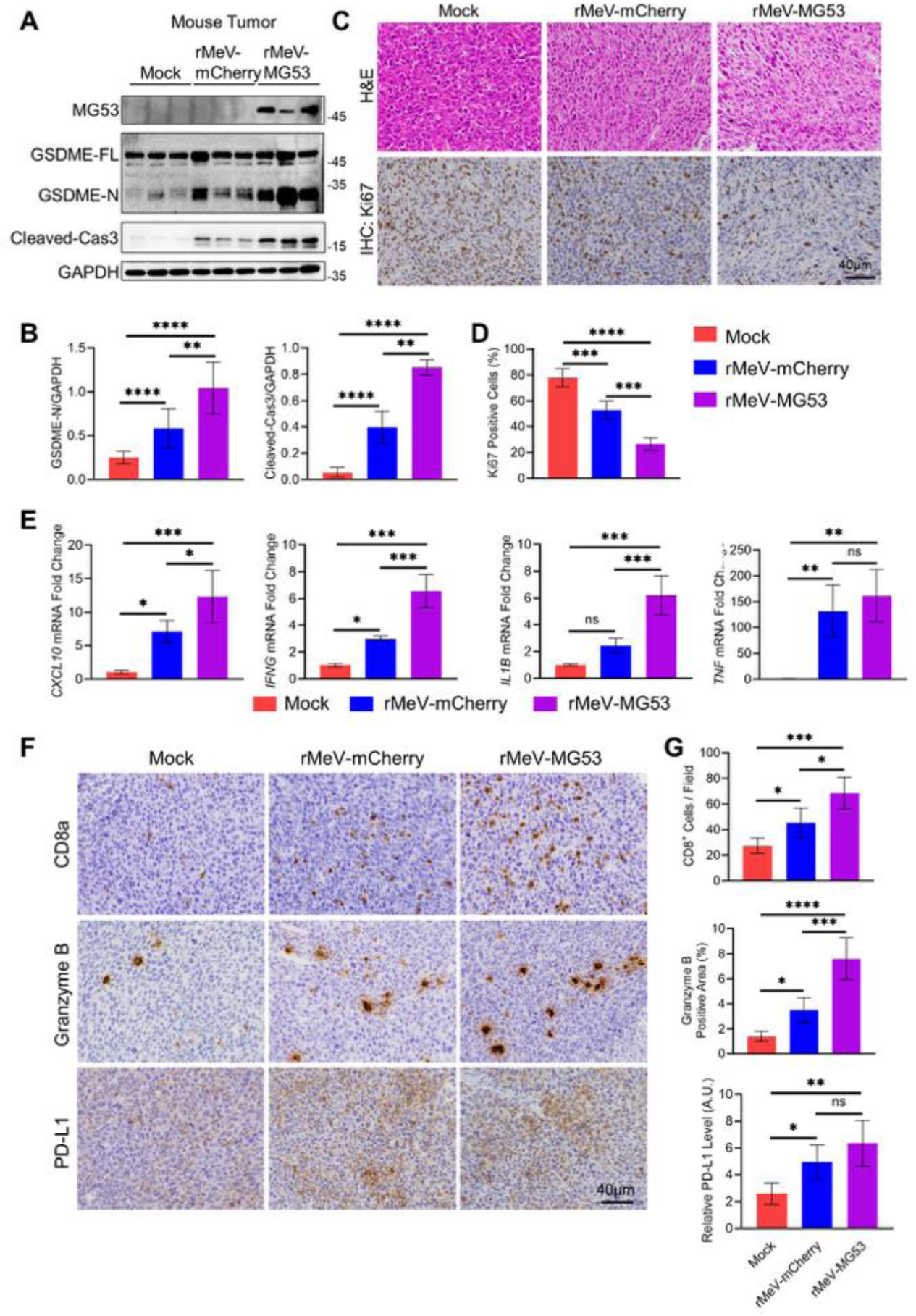
rMeV-MG53 drives *in vivo* pyroptosis, suppresses tumor proliferation, and remodels the tumor immune microenvironment. **(A)** Immunoblot analysis of MG53, GSDME-FL, GSDME-N, and cleaved caspase-3 in LL/2-hCD46 tumor lysates from mice treated with mock, rMeV-mCherry, or rMeV-MG53. GAPDH served as a loading control. **(B)** Densitometric quantification of GSDME-N and cleaved caspase-3 protein levels normalized to GAPDH, Data are presented as mean ± SD (*n*=3 per group). Representative H&E and Ki67 immunohistochemical staining of LL/2-hCD46 tumors from mock, rMeV-mCherry, and rMeV-MG53 groups **(C)**, with corresponding quantification of the percentage of Ki67⁺ cells **(D).** Scale bar, 40 µm (*n*=5 per group). **(E)** RT-qPCR analysis of *Cxcl10*, *Ifng*, *Il1b*, and *Tnf* expression in tumor tissues from the indicated treatment groups. Gene expression levels were normalized to the internal reference gene *Actb* and expressed as fold change relative to the mock group. Data are presented as mean ± SD (*n*=4 per group). **(F)** Representative immunohistochemical staining of CD8a, Granzyme B, and PD-L1. Scale bar, 40 µm. **(G)** Quantification of CD8a⁺ T-cells density per field (cells per field), Granzyme B– positive area (%), and relative PD-L1 expression level (A.U.) in tumors from the indicated treatment groups. Data are presented as mean ± SD (*n*=5 per group). Statistical significance was determined by one-way ANOVA followed by Tukey’s multiple-comparisons test. \**p* < 0.05; \*\**p* < 0.01; \*\*\**p* < 0.001; \*\*\*\**p* < 0.0001.

**Figure 6.**
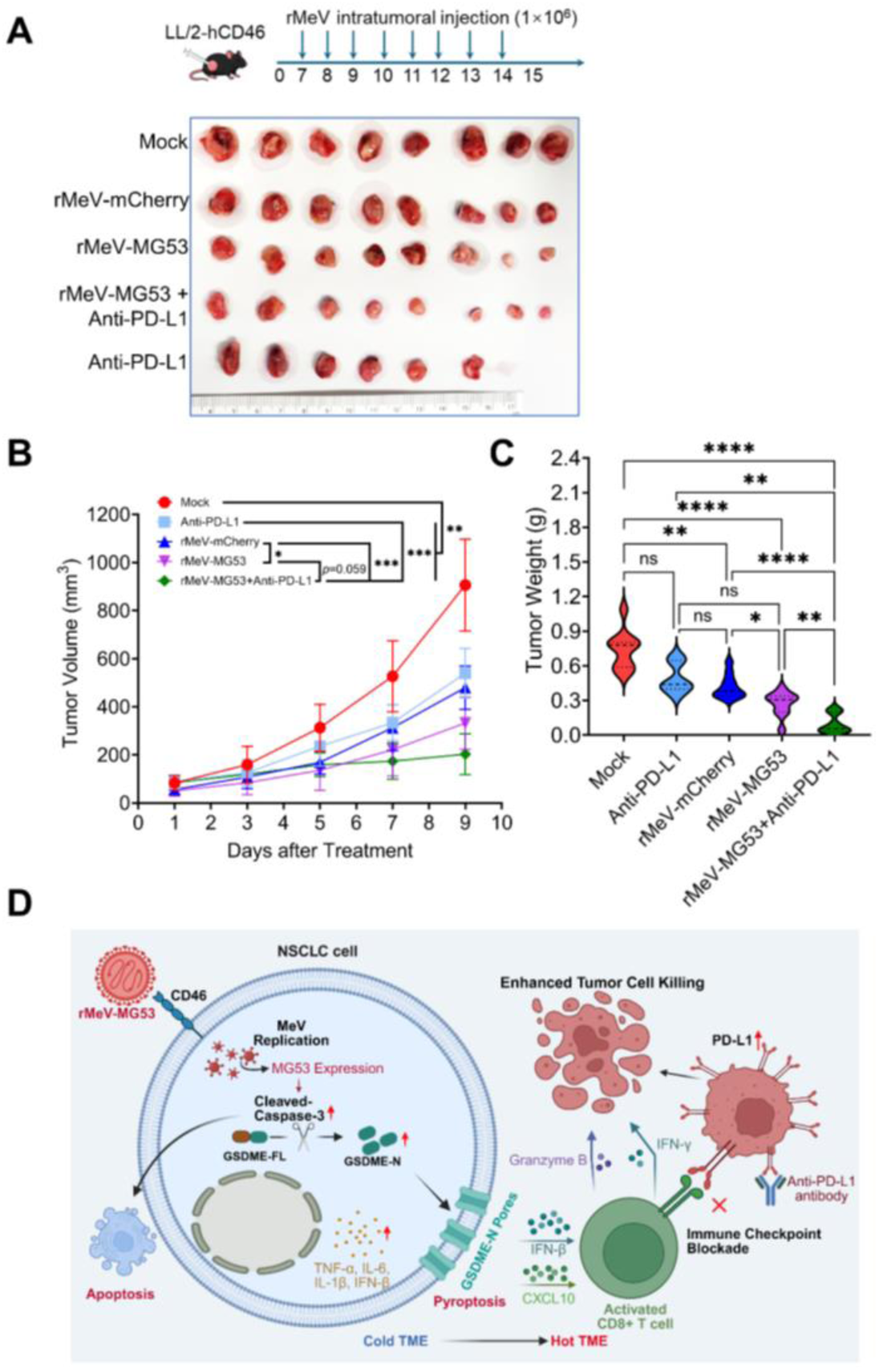
rMeV-MG53 combined with anti-PD-L1 blockade achieves superior tumor suppression in an immunocompetent syngeneic NSCLC model. (**A)** Treatment schematic and representative excised tumors from the five treatment groups cohort. Mice bearing LL/2-hCD46 tumors were treated with daily intratumoral viral injections (1 × 10⁶ PFU in 100 µL per dose) from days 7 through 14, with anti-PD-L1 antibody co-administered intratumorally in the combination group. **(B)** Longitudinal tumor volume growth curves for all five treatment groups. **(C)** Endpoint tumor weights presented as violin plots. Data are presented as mean ± SD (*n* = 6 for anti-PD-L1 group; n = 10 for all other groups). Tumor growth curves were analyzed by two-way repeated-measures ANOVA with Geisser–Greenhouse correction and Tukey’s multiple-comparisons test. Endpoint tumor weight was analyzed by Brown–Forsythe and Welch ANOVA with Dunnett’s T3 post hoc test. \**p* < 0.05; \*\**p* < 0.01; \*\*\**p* < 0.001; \*\*\*\**p* < 0.0001; ns, not significant. **(D)** Working model of rMeV-MG53 antitumor mechanism and synergy with PD-L1 blockade in NSCLC. rMeV-MG53 infects NSCLC cells *via* hCD46, driving viral replication and sustained MG53 expression. MG53 amplifies caspase-3 cleavage, triggering dual activation of apoptosis and GSDME-N–mediated pyroptosis, releasing pro-inflammatory mediators -including *TNF-α, IL-6, IL-1β, IFN-β*-that convert an immunologically cold TME into an immune-infiltrated hot TME. This drives *CXCL10*-mediated CD8⁺ T-cell recruitment and Granzyme B/IFN-γ– dependent cytotoxic tumor killing. Compensatory PD-L1 upregulation limits single-agent efficacy; combining rMeV-MG53 with anti-PD-L1 antibody relieves this adaptive checkpoint, unleashing full CD8⁺ T-cell cytotoxicity and enhanced antitumor control.

Consistent with this enhanced tumor-cell killing, Ki67 immunohistochemistry revealed a progressive decline in proliferating cells from mock to rMeV-mCherry to rMeV-MG53, with the rMeV-MG53 group showing significantly lower proliferation than both mock (*p* < 0.0001) and rMeV-mCherry (*p* < 0.001) (**Figure 5C and D**).

Next, we performed RT-qPCR profiling of the tumor tissue to understand the cytokine transcripts. rMeV-MG53 induced the most potent upregulation of *Cxcl10*, *Ifng*, and *Il1b*. Specifically, *Cxcl10* and *Ifng* transcripts were significantly elevated by rMeV-MG53 compared to both mock (*Cxcl10*: *p* < 0.001; *Ifng*: *p* < 0.001) and rMeV-mCherry (*Cxcl10*: *p* < 0.05; *Ifng*: *p* < 0.001). While *Il1b* induction by rMeV-mCherry was not statistically significant relative to mock, rMeV-MG53 drove a profound increase in *Il1b* over both mock (*p* < 0.001) and rMeV-mCherry (*p* < 0.001). In contrast, *TNF* was significantly upregulated by both viruses (*p* < 0.01 vs. mock), but no difference was observed between the two viral groups. This indicates that MG53-mediated cytokine amplification is selective —predominantly enhancing *Cxcl10*, *Ifng*, and *Il1b*—rather than representing a global transcriptional surge (**Figure 5E**).

This pronounced chemokine and interferon signature suggested enhanced recruitment of cytotoxic lymphocytes, which was subsequently verified *via* IHC. The density of CD8a⁺ T cells and the Granzyme B-positive area increased progressively across the mock, rMeV-mCherry, and rMeV-MG53 groups. rMeV-MG53 achieved the highest infiltration metrics for both markers (CD8a: *p* < 0.05 for mock vs. rMeV-mCherry, *p* < 0.0001 for mock vs. rMeV-MG53, *p* < 0.01 for rMeV-mCherry vs. rMeV-MG53; Granzyme B: *p* < 0.05, *p* < 0.0001, and *p* < 0.001, respectively). These findings demonstrate that MG53-armed virotherapy drives effector T-cell infiltration into the tumor TME (**Figure 5F and G**).

In parallel, we observed a compensatory upregulation of the PD-L1 immune checkpoint. Tumor PD-L1 protein levels rose significantly following either viral treatment compared to mock (mock vs. rMeV-mCherry, *p* < 0.05; mock vs. rMeV-MG53, *p* < 0.01) but did not differ between the two virus groups (**Figure 5F and G**). Therefore, PD-L1 upregulation represents a general adaptive response to oncolytic MeV infection rather than an MG53-specific effect. Recognizing that this adaptive resistance could attenuate the T-cell response elicited by the MG53 armed virus, we reasoned that combining rMeV-MG53 with PD-L1 blockade might translate this immune infiltration into improved tumor control.

### rMeV-MG53 combined with anti-PD-L1 therapy improves outcomes in an immunocompetent lung cancer model

To test this, tumor-bearing mice were divided into five groups: mock, anti-PD-L1 alone, rMeV-mCherry, rMeV-MG53, and rMeV-MG53 + anti-PD-L1. Recombinant virus (1 × 10⁶ PFU in 100 µL per dose) and anti-PD-L1 antibody were co-administered intratumorally daily from day 7 through day 14 (**Figure 6A**). At endpoint, excised tumors were smallest in the rMeV-MG53 + anti-PD-L1 group (**Figure 6A**). Tumor growth curves separated by treatment intensity: Mock grew fastest; anti-PD-L1 and rMeV-mCherry produced only modest inhibition; rMeV-MG53 alone was more effective; and the combination achieved the strongest suppression (**Figure 6B**). This combination was significantly superior to mock (*p < 0.01*) and to rMeV-mCherry (*p < 0.001*), and showed a trend toward greater suppression than rMeV-MG53 monotherapy although they did not reach statistical significance (*p* = 0.059). Furthermore, rMeV-MG53 alone remained superior to rMeV-mCherry (*p* < 0.05) (**Figure 6B**). The final tumor weights validated the observed differences in growth kinetics. The combination therapy resulted in the lowest overall tumor burden, significantly lighter than mock (*p* < 0.0001), anti-PD-L1 (*p* < 0.01), rMeV-mCherry (*p* < 0.0001), and rMeV-MG53 alone (*p* < 0.01). rMeV-MG53 monotherapy was also significantly lighter than mock (*p* < 0.0001) and rMeV-mCherry (*p* < 0.05) (**Figure 6C**). As expected, anti-PD-L1 alone did not significantly alter tumor weight compared to mock, reflecting the limited efficacy of checkpoint inhibitors as single agents in this model.

## Discussion

In this study, we demonstrate that engineering oncolytic MeV with the tumor suppressor MG53 (rMeV-MG53) enhances both direct cytotoxicity and antitumor immunity in NSCLC, converging on activation of the caspase - 3/GSDME pyroptotic axis. rMeV-MG53 preserved the replication competence of the parental MeV vector while significantly exceeding an unarmed control in killing NSCLC cells, amplified pro-inflammatory cytokine signaling both intrinsically and during infection, drove robust effector T-cell infiltration *in vivo*, and sensitized tumors to PD-L1 blockade. Together, these findings establish MG53 as a mechanistically distinct and therapeutically attractive payload for oncolytic virus engineering for the treatment of lung cancer.

The attenuated MeV Edmonston vaccine strain has been extensively investigated as an oncolytic platform across multiple cancer types, with favorable safety profiles documented in early-phase clinical trials(25, 29, 30, 40). The tractable MeV reverse genetics system enables insertion of therapeutic transgenes as additional transcription units, a strategy that has been exploited to arm MeV with a range of immunomodulatory payloads including GM-CSF, IL-12, anti-PD-L1 antibody fragments, and sodium iodide symporter(25, 30, 40). To our knowledge, this is the first study to employ MG53 as a therapeutic transgene in an oncolytic virus platform, thereby expanding the repertoire of therapeutic payloads available for armed oncolytic virotherapy. MeV infects cells using CD150/SLAM, nectin-4, and the complement regulator CD46, the last of which is commonly overexpressed on the surface of tumor cells as a mechanism to evade complement-mediated lysis(29, 41). However, the murine orthologs of these receptors do not support efficient MeV entry, precluding viral replication in murine tumor cells. To circumvent receptor restriction barriers, we engineered murine cells with stable expression of hCD46, establishing a syngeneic, MeV-permissive model that—unlike immunodeficient xenografts—permits comprehensive assessment of virus-induced immune responses and their interaction with checkpoint blockade in an immunocompetent host. Importantly, the viral delivery strategy employed here directly addresses a critical pharmacokinetic limitation of systemically administered recombinant MG53 protein, which exhibits a circulating half-life of approximately 1.5 h in preclinical models(10, 14). By encoding MG53 within the replicating viral genome, rMeV-MG53 achieves continuous, tumor-localized MG53 production throughout the course of viral replication, enabling sustained intratumoral MG53 exposure that is fundamentally unattainable through conventional protein-based delivery. This pharmacokinetic rationale represents a key translational advantage of the rMeV-MG53 platform over direct rhMG53 administration.

A central finding of this work is that MG53 potentiates GSDME-mediated pyroptosis rather than simply augmenting apoptosis. While oncolytic viruses, including MeV, have long been appreciated for their capacity to induce immunogenic cell death through multiple pathways(42), the specific contribution of gasdermin-mediated pyroptosis to oncolytic virotherapy efficacy has only recently gained attention(31, 43). Our data show that rMeV-mCherry alone was sufficient to induce detectable caspase-3 activation and GSDME cleavage, consistent with reports that MeV infection engages apoptotic and pyroptotic machinery as part of its native cytopathic program(31, 43). However, MG53 arming substantially amplified this response, suggesting that MG53 acts upstream or in parallel with viral-intrinsic death signaling to lower the threshold for pyroptotic execution. Critically, the pan-caspase inhibitor Z-VAD-FMK, but not the necroptosis inhibitor Necrostatin-1 or the ferroptosis inhibitor Ferrostatin-1, rescued cell viability following rMeV-MG53 infection, supporting a model in which caspase-3 serves as the convergence point for both apoptotic and pyroptotic death following infection, consistent with established GSDME biology in which caspase-3 cleavage acts as a switch between non-lytic apoptosis and lytic pyroptosis depending on GSDME expression level(32, 33). The mechanism by which MG53 potentiates this axis remains to be fully defined; given MG53’s known functions as an E3 ubiquitin ligase and membrane-associated protein(9, 22). It is plausible that MG53 modulates GSDME stability, caspase-3 activity, or membrane dynamics required for pore formation, and this represents an important direction for future mechanistic study.

A second key finding is that MG53 is not merely a cytotoxic payload but an intrinsically immunomodulatory one. Transcriptomic profiling of MG53-overexpressing cells in the absence of virus revealed enrichment of TNF, NF-κB, and IL-17 signaling, indicating that MG53 primes a pro-inflammatory cellular state independent of infection. This intrinsic property translated into amplified type I interferon and pro-inflammatory cytokine induction during active infection *in vitro*, and into a more selective but pronounced chemokine and cytokine signature *in vivo*, with rMeV-MG53 preferentially upregulating *Cxcl10, Ifng,* and *Il1b* without a global, non-specific transcriptional surge. *CXCL10* in particular is a well-established chemoattractant for CXCR3⁺ effector T cells(44), and its selective induction likely contributes mechanistically to the enhanced CD8⁺ T-cell and Granzyme B⁺ infiltration we observed in rMeV-MG53–treated tumors. Our RT-qPCR data, which demonstrate marked upregulation of pro-inflammatory cytokines (*IFNB1*, *IL6*, *CXCL10*, *IFNG*, *TNF*, and *IL1B*) in rMeV-MG53–infected cells and tumors, provide direct molecular evidence of this TME reprogramming. Together with the pyroptosis data, these findings support a model in which MG53 arming engages two complementary, mutually reinforcing mechanisms—lytic, DAMP-releasing pyroptotic cell death and chemokine-driven lymphocyte recruitment—to convert tumors from an immunologically quiescent state toward one more conducive to checkpoint immunotherapy.

The superior antitumor activity observed with rMeV-MG53 plus anti-PD-L1 combination therapy in the immunocompetent LL/2-hCD46 syngeneic model, provides compelling *in vivo* validation that pyroptosis-driven TME reprogramming sensitizes lung tumors to immune checkpoint blockade. These findings are consistent with accumulating evidence that OV-induced immunogenic cell death synergizes with ICI across multiple cancer types(40), and position rMeV-MG53 as a particularly potent immunogenic cell death-inducing OV platform due to the dual apoptosis/pyroptosis mechanism of tumor cell killing. By converting an immunologically excluded tumor into an inflamed, T cell-infiltrated one, rMeV-MG53 creates a favorable context for anti-PD-L1 therapy to exert its full activity.

This immune remodeling, however, was accompanied by compensatory PD-L1 upregulation following viral treatment, consistent with the well-described phenomenon of adaptive immune resistance, whereby interferon-driven T-cell infiltration triggers PD-L1 induction as a tumor-intrinsic feedback mechanism(45). Notably, PD-L1 upregulation did not differ between rMeV-mCherry and rMeV-MG53 groups despite their markedly different degrees of T-cell infiltration, suggesting that PD-L1 induction in this model may be driven largely by general interferon signaling associated with viral infection rather than scaling proportionally with the magnitude of immune infiltration. This finding reinforces the rationale for combining rMeV-MG53 with PD-L1 blockade, as it indicates that even modest oncolytic-virus-induced interferon signaling can establish an adaptive resistance checkpoint that single-agent virotherapy alone is unlikely to overcome. Indeed, the combination of rMeV-MG53 with anti-PD-L1 antibody achieved the strongest tumor control among all treatment groups, while anti-PD-L1 monotherapy showed minimal efficacy—consistent with the limited single-agent activity of checkpoint inhibitors in poorly infiltrated NSCLC and underscoring the value of oncolytic priming to unlock checkpoint blockade responsiveness(46).

MG53’s E3 ubiquitin ligase activity has been shown to suppress tumor growth through ubiquitination-dependent degradation of multiple oncogenic substrates, including G3BP2, RAC1, and cyclin D1 across diverse cancer types including NSCLC, hepatocellular carcinoma, and colorectal cancer(20, 22, 47). In the context of rMeV-MG53, intratumoral MG53 expression is anticipated to simultaneously suppress oncogenic signaling pathways, enhance apoptotic sensitivity, and amplify the caspase-3/GSDME pyroptotic axis — a multifaceted antitumor activity that is mechanistically complementary to, and synergistic with, MeV-mediated direct oncolysis. The observation that rMeV-MG53 consistently outperforms parental rMeV in cytotoxicity assays across multiple NSCLC cell lines supports the conclusion that MG53 transgene expression provides meaningful additive antitumor activity beyond viral oncolysis alone.

Several limitations of this study merit mention: First, as MeV is a primate-adapted pathogen with limited replication in immunocompetent rodents, the LL/2 syngeneic model used here does not fully recapitulate human MeV replication kinetics and spread. This is a well-recognized challenge in the preclinical development of oncolytic MeV and argues for future validation in hCD46-transgenic mouse models or non-human primate systems that better approximate MeV biology in humans(48). Alternatively, a comparative medicine approach using canine pulmonary adenocarcinoma could be considered. Canine pulmonary adenocarcinoma has been proposed as a model for NSCLC(49). Studies have also demonstrated that MeV lacking its binding affinity for SLAM has an antitumor effect on canine lung cancer cell line(50). Translation of our work to canine pulmonary adenocarcinoma may facilitate investigation in an immunocompetent spontaneous disease model that may more accurately reflect NSCLC. Second, while our data demonstrate GSDME-N generation and morphological features of pyroptosis *in vitro*, direct quantification of pyroptosis-associated DAMP release - including HMGB1, ATP, and IL-1β —and genetic validation using GSDME-deficient cells would further strengthen the mechanistic link between GSDME-mediated pyroptosis and the enhanced antitumor activity of rMeV-MG53. In addition, although increased CD8⁺ T-cell infiltration and granzyme B expression support enhanced antitumor immunity, comprehensive profiling of the tumor immune microenvironment, including macrophages, NK cells, dendritic cells, regulatory T cells, and myeloid-derived suppressor cells in tumors following rMeV-MG53 treatment *in vivo* would further strengthen the mechanistic link between pyroptotic cell death and the observed anti-PD-L1 synergy. These analyses represent important objectives for future studies. Third, the potential contribution of MG53-mediated E3 ligase activity to the enhanced caspase-3/GSDME signaling observed requires further mechanistic dissection, including assessment of key MG53 ubiquitination substrates in the context of viral infection. Finally, the optimal dosing regimen, scheduling, and route of administration for rMeV-MG53 and anti-PD-L1 combination therapy will require systematic optimization in future preclinical studies prior to clinical translation.

Based on our collective findings, we propose the following working model (**Figure 6D**). Upon intratumoral delivery, rMeV-MG53 infects NSCLC cells *via* hCD46 or other related receptors, achieving sustained MG53 expression through viral replication. Virus-delivered MG53 amplifies caspase-3 cleavage, triggering dual apoptosis and GSDME-mediated pyroptosis, driving release of *TNF-α, IL-6, IL-1β*, and *IFN-β* and converting an immunologically “cold” TME into an immune-infiltrated “hot” TME. *CXCL10* induction recruits CD8⁺ effector T cells, which exert cytotoxicity *via* Granzyme B and IFN-γ. However, this immune activation simultaneously induces compensatory PD-L1 upregulation on tumor cells, establishing an adaptive resistance checkpoint that constrains single-agent virotherapy efficacy. Combining rMeV-MG53 with anti-PD-L1 blockade relieves this checkpoint, fully unleashing infiltrating CD8⁺ T-cell cytotoxicity and achieving the strongest antitumor control observed across all treatment conditions. This working model positions rMeV-MG53 + anti-PD-L1 as a mechanistically rational combination strategy to sensitize immune-excluded NSCLC to checkpoint immunotherapy.

In conclusion, this study establishes MG53 as a rationally designed payload for oncolytic MeV that simultaneously enhances direct tumor cytotoxicity through GSDME-mediated pyroptosis and remodels the tumor immune microenvironment to favor effector T-cell infiltration. By coupling this dual mechanism with PD-L1 blockade, rMeV-MG53 achieved superior tumor control in an immunocompetent NSCLC model, offering a promising combinatorial strategy to sensitize immunologically cold or checkpoint-resistant lung tumors to immunotherapy. These findings provide a strong mechanistic and preclinical foundation for the further development of rMeV-MG53 as a novel therapeutic strategy for lung cancer.

## Acknowledgements

This research project has been supported by the American Lung Association Innovation Award (IA-1613730); The Ohio State University Comprehensive Cancer Center Startup Fund, Internal funds from the Ohio State University: Infectious Diseases Institute (IDI) Interdisciplinary Seed Fund, and Integrated Oncology Fund to H.L. This research is also partially supported by an NIH/NIAID award (1R01AI183671-01A1) and an NIH/NCI award (5U01CA294881) to J. L. This study used the Shared Resources and Core Facilities at The Ohio State University Comprehensive Cancer Center for Microscopy, Flow Cytometry, and Pathology and Imaging that are supported in part by the NIH CCSG P30 grant (P30CA016058).

## Author contributions

H.L., J. L., and Z. L. designed the studies. Z.L., C.H., F.J., M.B., U.L., S.L., S. Z., P.L., H. P., and X. L. conducted experiments and data analysis. Z.L., J.L., and H.L. prepared and revised the manuscript. All authors have read and agreed to the published version of the manuscript.

## Data Availability

The datasets for this study are available from the corresponding author on reasonable requests.

## Artificial Intelligence Tools Use Statement

The authors utilized Claude (Anthropic, San Francisco, CA, USA), a large language model AI assistant, to assist with grammar checking, proofreading, and language editing of this manuscript. All scientific content, data, interpretation, and conclusions were generated and verified solely by the authors.

